# Fast tumor phylogeny regression via tree-structured dual dynamic programming

**DOI:** 10.1101/2025.01.24.634761

**Authors:** Henri Schmidt, Yuanyuan Qi, Benjamin J. Raphael, Mohammed El-Kebir

## Abstract

**Motivation:** Reconstructing the evolutionary history of tumors from bulk DNA sequencing of multiple tissue samples remains a challenging computational problem, requiring simultaneous deconvolution of the tumor tissue and inference of its evolutionary history. Recently, phylogenetic reconstruction methods have made significant progress by breaking the reconstruction problem into two parts: a regression problem over a fixed topology and a search over tree space. While effective techniques have been developed for the latter search problem, the regression problem remains a bottleneck in both method design and implementation due to the lack of fast, specialized algorithms.

**Results:** Here, we introduce *fastppm*, a fast tool to solve the regression problem via *tree-structured dual dynamic programming. fastppm* supports arbitrary, separable convex loss functions including the *ℓ*_2_, piecewise linear, binomial and beta-binomial loss and provides asymptotic improvements for the *ℓ*_2_ and piecewise linear loss over existing algorithms. We find that *fastppm* empirically outperforms both specialized and general purpose regression algorithms, obtaining 50-450*×* speedups while providing as accurate solutions as existing approaches.

Incorporating *fastppm* into several phylogeny inference algorithms immediately yields up to 400*×* speedups, requiring only a small change to the program code of existing software. Finally, *fastppm* enables analysis of low-coverage bulk DNA sequencing data on both simulated data and in a patient-derived mouse model of colorectal cancer, outperforming state-of-the-art phylogeny inference algorithms in terms of both accuracy and runtime.

**Availability:** *fastppm* is implemented in C**++** and available as both a command-line interface and Python library at github.com/elkebir-group/fastppm.git.

## Introduction

Inferring the evolutionary history of tumors, known as *phylogenetic reconstruction*, is a fundamental challenge in cancer genomics. Over the past two decades, phylogenetic reconstruction of tumors has been performed using a variety of sequencing technologies, including bulk DNA sequencing [1], single-cell DNA sequencing [2–4], single-cell RNA sequencing [5], and lineage tracing [6, 7] technologies. Concurrently, reconstruction has been performed using a variety of phylogenetic markers, such as somatic single-nucleotide variants (SNVs) [8–14], copy number aberrations [15, 16], gene expression [5], and DNA methylation [17]. Despite the diversity of sequencing technologies and markers used to perform phylogenetic reconstruction of tumors, reconstructing tumor phylogenies from bulk DNA sequencing of tumor tissue remains important, as large-scale cohort studies of patient tumors [1, 18, 19] continue to apply the technology. A key difficulty in phylogenetic reconstruction of tumors from bulk DNA sequencing data is that sequencing measures a *mixture* of the underlying clonal genotypes. Effective methods for phylogenetic reconstruction [11, 12, 20, 21, 13, 14, 22] thus must simultaneously deconvolve the mixtures while performing tree reconstruction. However, it has recently been observed [23] that this phylogenetic reconstruction problem is highly under-determined, leading to non-uniqueness of solutions, i.e. the presence of multiple tumor phylogenies that explain the data equally well. Further, many tumor phylogeny methods approximate the raw read counts with estimated frequencies in order to speed up phylogenetic inference, leading to a loss of signal and accuracy. Finally, to manage computational intractability, many existing analyses and methods perform reconstruction on *clusters* of mutations [24], but this fails to leverage useful phylogenetic signal and relies on a correct clustering.

Envisioning that the next-generation of methods for phylogenetic reconstruction from bulk DNA sequencing will account for non-uniqueness of the solution space, accurately model read counts, and attempt reconstruction directly from mutation-level counts, we sought to build a framework promoting these developments. The key computational bottleneck hindering such development is the lack of specialized algorithms for a *regression* problem over a fixed topology. Specifically, all scalable methods [13, 14, 22] for phylogenetic reconstruction work by breaking the reconstruction problem into two parts, a repeatedly-solved Perfect Phylogeny Regression (PPR) problem over a fixed topology and a search over tree space. In the PPR problem, one seeks latent mutation frequencies **f** for a fixed tree topology 𝒯 best explaining the observed read counts **v, d**, quantified via a specified loss function *L*(**f**) (Fig. 1a,b). Current methods to solve the PRR problem either employ general-purpose convex optimization packages or specialized solvers (Table 1). However, general-purpose convex optimization packages do not leverage the unique tree structure inherent to the PPR problem. On the other hand, solvers that leverage the tree structure have only been developed for restrictive loss functions *L*(**f**) operating directly on frequencies and ignoring additional information present in the read counts, such as the *ℓ*_1_ [22] and *ℓ*_2_ loss [25].

**Table 1.**
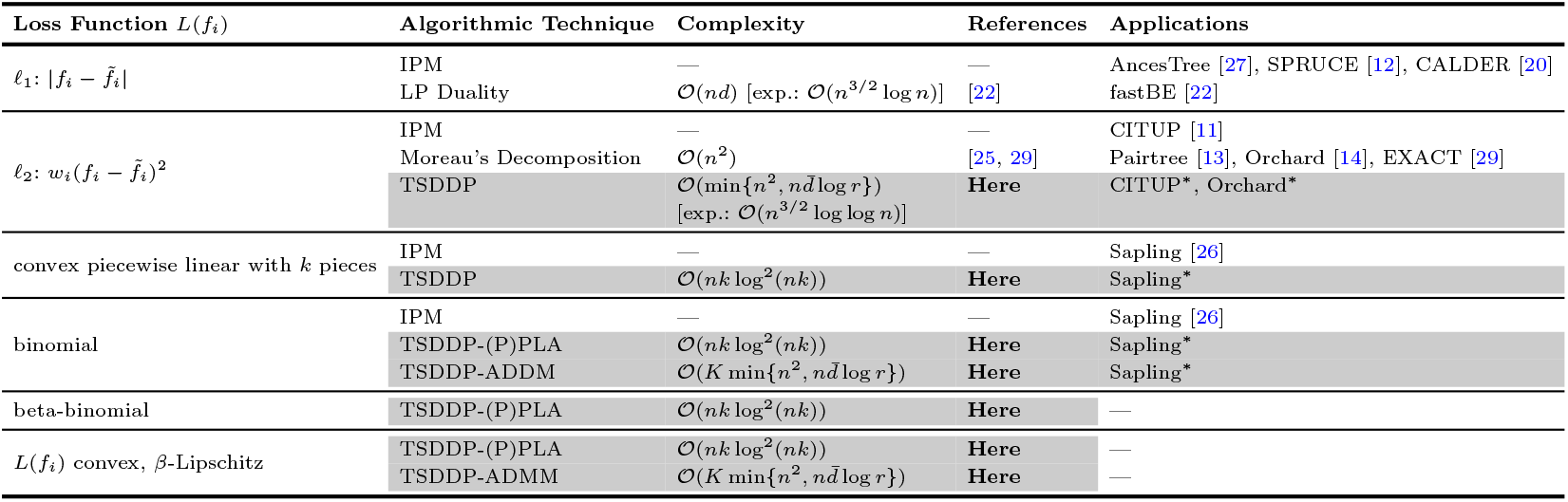
Properties of algorithms for solving the Perfect Phylogeny Regression problem with a loss function of the form 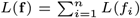 where *n* is the size of the clonal tree 𝒯, *d* is the maximum depth of 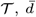 is the average depth of 𝒯, *r* is the maximum number of children in 𝒯, *k* is the parameter used in the *k*-PLA and PPLA(*k, τ, σ*) algorithms, and *K* is the number of iterations used in the alternating directions method of multipliers (ADMM) algorithm. TSDDP refers to the tree structured dual dynamic programming technique introduced in this work. Moreau’s decomposition refers to the method for constructing the dual developed in [25, 29]. IPM refers to interior point methods which form the basis of state-of-the-art convex optimization software. The notation [exp.:] refers to the expected runtime over random classes of *n*-vertex trees [30].

**Fig. 1.**
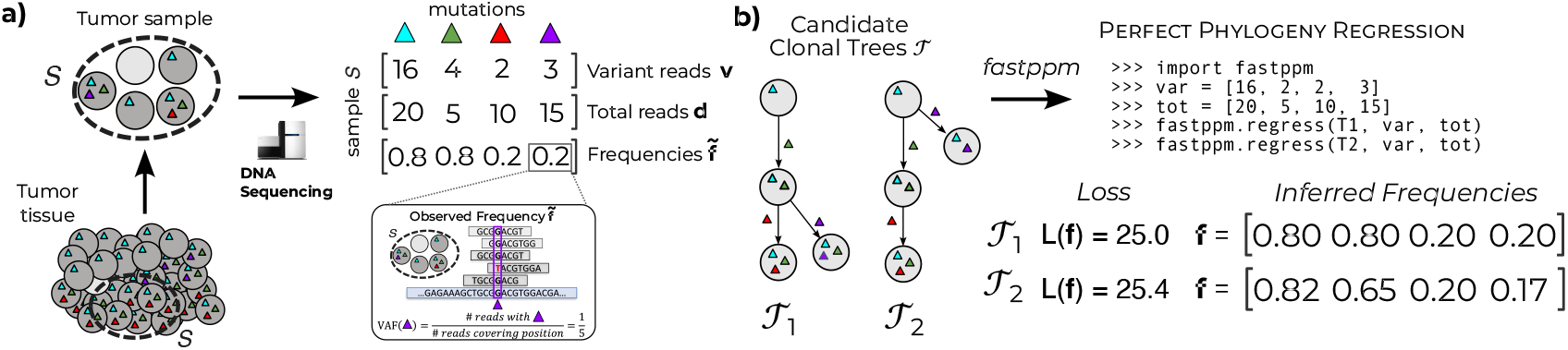
**a)** Variant read counts **v** and total read counts **d** are obtained through bulk DNA sequencing of tumor tissue followed by sequence alignment and variant calling. **b)** Provided a clonal tree *𝒯*, variant read counts **v**, and total read counts **d**, *fastppm* solves the Perfect Phylogeny Regression problem to obtain the minimum loss along with inferred frequencies **f**, enabling the ranking of candidate clonal trees.

To overcome the key computational bottleneck in tumor phylogeny reconstruction from bulk DNA sequencing data, we developed *fastppm. fastppm* solves the Perfect Phylogeny Regression problem via tree-structured dual dynamic programming (TSDDP), a new technique that supports arbitrary loss functions of the form 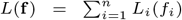 where *L*_*i*_ is convex. This includes previously studied loss functions such as the *ℓ*_1_ and (weighted) *ℓ*_2_ loss, for which we obtain *asymptotic* improvements over existing algorithms [25, 22], along with loss functions modeling commonly used probability distributions for read count data such as the binomial and beta-binomial distributions. We implemented *fastppm* in C**++**, and made it available as both a shared library and via a command-line interface. Moreover, *fastppm* provides Python bindings, making *fastppm* easy-to-use in existing tumor phylogeny inference pipelines (Fig. 1b). We demonstrate the advantages of *fastppm* on simulated tumors, showing speedups of 100*×* for the *ℓ*_2_ loss and 400*×* for the binomial loss. Incorporating *fastppm* into state-of-the-art tumor phylogeny methods such as CITUP [11], Orchard [14], and Sapling [26] consequently led to significant speed-ups of 3*×* to 400*×*. Specifically, *fastppm* enables Sapling to scale beyond 500 mutations, while the original implementation could only be run up to 50 mutations. On real data, we show that *fastppm* enables one to use accurate loss functions that operate on read counts rather than frequencies. This in turn enables more accurate tree inference, especially in a shallow coverage setting where one might opt to trade-off coverage for sequencing more regions/biopsies of a tumor.

### Problem Statement

We consider the problem of tumor phylogeny inference from bulk DNA sequencing data. After aligning sequencing reads to the reference genome and performing variant calling (Fig. 1a), we obtain the variant and total read count matrices **V, D** *∈* ℕ ^*m×n*^ of *n* single-nucleotide variants (SNVs) across *m* samples. In this manuscript, we will refer to SNVs as mutations. For each mutation *i* in sample *p*, we denote the number of variant and total reads by *v*_*pi*_ and *d*_*pi*_, respectively. In addition, the quantity 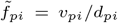 is the *observed frequency*, also known as the variant allele frequency of mutation *i* in sample *p*.

The *n* mutations of the tumor evolve down a rooted tree, called a *clonal tree* 𝒯 [27, 22], such that each mutation is gained exactly once and never subsequently lost — this is known as the infinite sites assumption (ISA) [28]. We note that, while the infinite sites assumption assumption may be violated for individual mutations due to copy-number loss, tumor phylogeny inference pipelines correct for such events by inferring mutation clusters for which the ISA holds [24, 1]. We will consider mutation clusters as individual mutations^1^.

The *n* vertices or *clones* of 𝒯 are uniquely identified by the mutation on the edge into the clone, and are labeled as [*n*] = {1, …, *n*}. Associated with each clone *i* of 𝒯 is a *n*-dimensional binary vector **b**_**i**_ *∈* {0, 1}^*n*^ recording the mutations present in clone *i*. Specifically, *b*_*ij*_ = 1 if and only if mutation *j* is present in clone *i*. Given a clonal tree 𝒯, we denote its root node as *r* = |𝒯), its vertex set as *V* (𝒯), and its edge set as *E*(𝒯). Further, the parent (if it exists) of vertex *i* in *T* is denoted as *π*(*i*), its children as *δ*(*i*), and its descendants (including itself) as *D*(*i*).

As bulk DNA sequencing measures a mixture of clones, we denote with *u*_*pi*_ ≥ 0 the proportion of clone *i* in sample *p*. Consequently, the underlying frequency *f*_*pi*_ with which we observe mutation *i* in sample *p* is 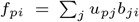.Compactly, this relation can be written in matrix form as **F** = **UB** where **B** = [*b*_*ij*_] is the *n*-by-*n clonal matrix* associated with *T*, **F** = [*f*_*pi*_] is the *m*-by-*n frequency matrix*, and **U** = [*u*_*pi*_] is the *m*-by-*n usage matrix*. There is a one-to-one correspondence between clonal matrices **B** and clonal trees *T* [27, 22]. Note that that the underlying frequency *f*_*pi*_ might not be equal to the observed frequency 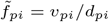 obtained from the observed variant and total read counts due to sampling uncertainty.

The goal in tumor phylogeny inference is to infer the true clonal tree 𝒯, the underlying frequencies **F**, and the usage matrix **U**, provided the read count data **V, D** *∈* ℕ^*m×n*^. To infer and/or summarize tumor phylogeny solution spaces, existing methods repeatedly solve the following optimization problem for either different clonal trees 𝒯 [11, 22, 29] or in a progressive fashion by growing clonal trees one mutation at a time [12, 20, 13, 14, 26]. Moreover, existing methods use different loss functions (summarized in Table 1).

#### Problem 1

(Perfect Phylogeny Regression (PPR)).

*Given variant and total read counts* **V, D** *∈* ℕ^*m×n*^ *for m samples and n mutations, a clonal matrix* **B** *∈* {0, 1} ^*n×n*^ *associated with a clonal tree* 𝒯 *and loss function L*(**F** | **V, D**) : [0, 1]^*m×n*^ → ℝ_≥0_, *find a frequency matrix* **F** *∈* [0, 1]^*m×n*^ *and usage matrix* **U** *∈* [0, 1]^*m×n*^, **U**𝟙≤ 1 *such that* **F** = **UB** *and L*(**F** | **V, D**) *is minimized*.

We denote the value of the minimizer as *L*^*^(𝒯). When the loss function separates across samples as 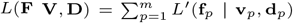, which is typical, one can solve the PPR problem independently for each sample. Thus, we assume *m* = 1 and use row vectors **f** ^*T*^ and **u**^*T*^ rather than the matrices **F** and **U**.

### Theory and Algorithms

Although specific cases of the PPR problem have been studied in the tumor phylogenetics literature, no general theory has developed for solving phylogeny constrained regression problems. By using Lagrangian duality as a starting point, we start to develop such a theory, showing that the PPR problem is equivalent to a *tree separable* optimization problem. Exploiting this tree separability, we derive an abstract dynamic programming algorithm for solving the PPR problem, an approach we call *tree structured dual dynamic programming (TSDDP)*. A summary of the algorithmic results obtained with TSDDP is provided by Table 1.

#### A tree separable dual problem

We generalize the dual construction used in [22] to derive an equivalent form of the PPR problem, which is *tree separable* in the coordinates of the dual variables, a term we define below. Rather than use linear programming (LP) duality, however, we employ the much stronger notion of Lagrangian duality [31]. Not only does this simplify the dual construction by allowing us to avoid an epigraphical reformulation of our problem, but it also allows us to extend the dual to arbitrary convex loss functions beyond the *ℓ*_1_ norm. As we will demonstrate in the next section, the tree separability of the dual problem allows for efficient optimization using a tree structured dynamic programming algorithm.

To start, fix a clonal matrix **B** and its associated clonal tree 𝒯, and assume the loss function *L* separates across mutations as 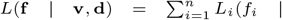 *v*_*i*_, *d*_*i*_). Note that this assumption typically follows directly from independence among the *n* mutations. Simplifying notation by omitting the variant and total read counts from the loss function, the PPR problem can be written mathematically as the following (primal) optimization problem,

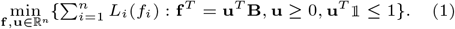

To simplify the preceding optimization problem, we invoke the basic theory of clonal matrices and trees developed in [27, 22], which states that for a given frequency vector **f** ^*T*^ there exists a usage vector **u**^*T*^ such that **f** ^*T*^ = **u**^*T*^ **B** if and only if **f** satisfies the following Sum Condition (SC),

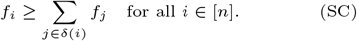

More formally, we use the following result, which can be proven both combinatorially [27] or algebraically [22].

##### Proposition 1.

*Let* 𝒯*be a clonal tree and* **B** *be its associated clonal matrix. Then, for a frequency vector* **f** ^*T*^ *there exists a usage vector* **u**^*T*^ *such that* **u** ≥ 0, **u**^*T*^ 𝟙≤ 1, *and* **f** ^*T*^ = **u**^*T*^ **B** *if and only if* **f** *satisfies the Sum Condition* (SC) *and has f*_*r*_ ≤ 1.

Applying Proposition 1 to the primal problem (1), we obtain the equivalent optimization problem which factors out the usage vector **u**:

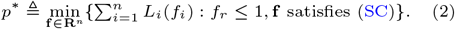

As **B** is invertible (in linear time [22]), we can recover the unique **u**^*^ corresponding to the minimizer **f** ^*^ in 𝒪(*n*) time. To derive the Lagrangian dual, we associate the dual vector 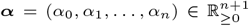 with the inequality constraints, obtaining the Lagrangian,

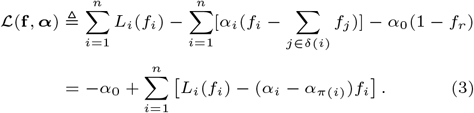

Here, the equality follows from the fact that each *f*_*i*_ appears twice in the right hand side of (3): once with *α*_*i*_ and once with the parent dual variable *α*_*π*(*i*)_. For simplicity, we are abusing notation and setting *π*(*r*) = 0.

The Lagrangian dual function *g*(***α***), then optimizes out the primal variable ***f***, yielding 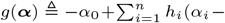 *α*_*π*(*i*)_) where *h*_*i*_(*x*) ≜ min_*f*_ {*L*_*i*_(*f*) − *xf*}. Importantly, the dual function *g*(***α***) is *tree separable* in the dual variable ***α***. We will see in the next section that the tree separability of *g*(***α***) enables us to solve the Lagrangian dual problem with tree structured dynamic programming. The Lagrangian dual problem is *d*^*^ ≜ max_***α***≥0_ *g*(***α***). Since ***α*** ≥ **0**, it is clear that *d*^*^ ≤ *p*^*^ for an arbitrary (not necessarily convex) set of loss functions *L*_*i*_. However, when *L*_*i*_ are convex, we obtain strong duality *d*^*^ = *p*^*^ as the constraint set is affine [31]. To complete this section, we summarize our results in the following theorem statement.

##### Theorem 1.

*Let* 𝒯 *be a clonal tree on n mutations. If the loss function separates as* 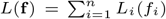 *with L*_*i*_ *all convex, then the PPR problem is equivalent to*

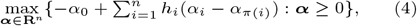

*where h*_*i*_(*x*) ≜ min_*f*_ {*L*_*i*_(*f*) − *xf*}.

#### Tree structured dual dynamic programming

We provide a meta algorithm for solving the tree separable dual problem using *tree structured dual dynamic programming (TSDDP)*. We describe the algorithm’s basic ingredients and several properties that are conserved across different choices of loss function.

To start, let *J*_*i*_ be the optimal solution to the PPR problem for the subtree rooted at node *i* when the dual variable of the parent *α*_*π*(*i*)_ is fixed. Formally, let

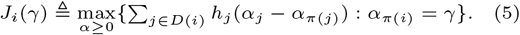

To solve the PPR problem it is sufficient to maximize over *J*_*r*_ since 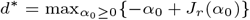 by definition. Due to the tree separability of the PPR problem, however, *J*_*r*_ is computable from *J*_*i*_ over the children *i ∈ δ*(*r*). Indeed, this holds recursively, yielding the following recursion:

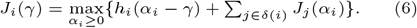

The above recurrence then implies the following bottom-up dynamic programming algorithm for computing the optimal value of PPR problem.

i. Fix a representation *ℛ* (*J*_*i*_) for each *J*_*i*_.
ii. Compute the representation *ℛ* (*J*_*i*_) at the leaf nodes.
iii. Compute the representation *ℛ* (*J*_*i*_) at a node *i* provided the representations *ℛ* (*J*_*j*_) at all children *j ∈ δ*(*i*).
iv. Solve the one-dimensional optimization problem 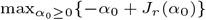 using the representation of the root node *ℛ* (*J*_*r*_).

To recover the optimal solution, we perform top-down backtracking over the tree topology, analogous to the backtracking performed to recover an optimal ancestral labeling in Sankoff’s algorithm [32]. In particular, we recover the dual variable 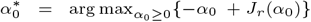.Then, given the optimal dual variable 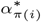 of the parent *π*(*i*) of *i*, we compute the optimal dual variable for node *i* as

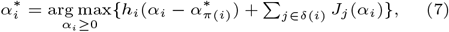

where we set *α*_*π*(*r*)_ = *α*_0_ for convenience. Since the representations of *J*_*j*_ have already been computed prior to backtracking, recovering the dual variables consists of solving *n* one-dimensional optimization problems. With the dual variables in hand, we finally recover the primal variables 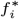 by computing 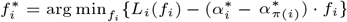.

In general, it is step (iii) of the TSDDP algorithm, the inductive step, which is the most challenging to implement, since one requires that the representation *ℛ* (*J*_*i*_) is closed under the recurrence relation. However, even in the general case, *J*_*i*_ possess several nice properties. In particular, *h*_*i*_ is the negation of the convex conjugate of *L*_*i*_, implying that *h*_*i*_ is concave, even when *L*_*i*_ is not necessarily convex (Observation 1). Next, it holds that *J*_*i*_ is always concave (Observation 2), implying that computing the one-dimensional maximizer of *J*_*i*_ is tractable.

#### Specializing TSDDP to *ℓ*_1_, *ℓ*_2_, and piecewise linear loss functions

Perhaps the simplest loss function is a quadratic, or *ℓ*_2_, loss. Indeed, one of the earliest methods for inferring tumor phylogenies from bulk DNA sequencing data, CITUP [11], used a quadratic loss function within a quadratic integer programming framework. The quadratic loss generalizes to probabilistic settings where the observed frequencies are drawn from a Gaussian distribution centered at the estimated frequencies [13, 14]. Jia et al. [25] solved the PPR problem under an unweighted *ℓ*_2_ loss with an 𝒪(*n*^2^) time algorithm by constructing a dual problem using Moreau’s decomposition and solving it iteratively.

With TSDDP, we derive an asymptotically faster algorithm for the weighted *ℓ*_2_ loss. To derive this algorithm, we fix our computational representation *ℛ* (*J*_*i*_) to be the value *J*_*i*_(0), along with the intercept, slopes, and breakpoints of the piecewise linear derivative 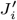. Computing the representation of the leaf nodes (ii) is easily done in 𝒪(1) time in closed form. The inductive step (iii) takes 𝒪(|*D*(*i*)| log |*δ*(*i*)|) time and the final one-dimensional optimization step (iv) takes 𝒪(*n*) time, leading to the desired time complexity using a careful analysis. The key technical challenge in obtaining our result is in proving that *J*_*i*_ has a piecewise linear derivative and that the representation of the derivative 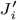 is quickly computed. Detailed derivations for each step of the algorithm are provided in Supplementary Results.

##### Theorem 2.

*TSDDP solves the PPR problem for the (weighted) ℓ*_2_ *loss* 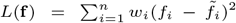 *in* 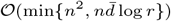 *time where* 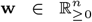 *are fixed weights*, 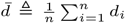 *is the average depth of a node in* 𝒯, *and* 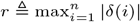 *is the maximum out-degree*.

Thus, we improve upon the 𝒪(*n*^2^) complexity of the Jia et al. [25] algorithm. Another key advantage of our algorithm is that *almost all* trees have 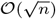 diameter and 𝒪(log *n*) maximum out-degree [30]. In addition, when trees are restricted to being binary, they typically have 𝒪(log *n*^2^) diameter [33]. Consequently, our algorithm for the *ℓ*_2_ loss has 𝒪(*n*^3*/*2^ log log *n*) expected complexity for random trees and 𝒪(*n* log *n*) expected complexity for random binary trees. The worst-case 𝒪(*n*^2^) runtime appears when 𝒯 is a path (i.e. 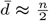).

Arguably, the next simplest loss function is the *ℓ*_1_ loss, which is found in several methods [27, 12, 20, 22] for inferring tumor phylogenies from bulk DNA sequencing data. To solve the PPR problem under an unweighted *ℓ*_1_ loss, [22] developed an algorithm running in 𝒪(*nd*) time, where *d* is the depth of 𝒯, by exploiting linear programming (LP) duality and the geometry of the *ℓ*_1_ loss.

Using TSDDP, we extend the case of an unweighted *ℓ*_1_ loss to convex piecewise linear loss functions, while also reducing the time complexity to 𝒪(*nk* log^2^(*nk*)). To derive this algorithm, we fix (i) the computational representation *ℛ* (*J*_*i*_) to be the slopes, breakpoints, and intercept of *J*_*i*_, which is shown to be a (concave) piecewise linear function. The key technical insight in obtaining the improved runtime is observing that the representations *ℛ* (*J*_*i*_) can be efficiently manipulated by storing the breakpoints and slopes in a specialized data structure which supports the addition and infimal conjugation of convex, piecewise linear functions [34]. Using such a data structure, steps (ii) and (iii) take an average of 𝒪(*k* log^2^(*nk*)) time. As the final one-dimensional optimization step (iv) consists of evaluating the representation *J*_*r*_ at all breakpoints, this leads the desired time complexity of 𝒪(*nk* log^2^(*nk*)). Detailed derivations are provided in Supplementary Results.

##### Theorem 3.

*TSDDP solves the PPR problem for a convex, piecewise linear loss* 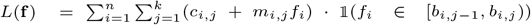 *in* 𝒪(*nk* log (*nk*)) *time*.

#### Extending TSDDP to arbitrary loss functions

A key advantage of solving the PPR problem for the convex quadratic and piecewise linear loss functions is that it enables approximation of non-linear (convex) loss functions. Here, we present two such approaches to solving PRR for arbitrary convex loss functions using *fastppm* for the special cases of the *ℓ*_2_ and piecewise linear loss functions as a subroutine. We then apply these algorithms to the binomial and beta-binomial loss functions.

##### A structural alternating directions method of multipliers approach

The alternating directions method of multipliers (ADMM) [35] reduces a constrained convex optimization to a sequence of *simpler* convex subproblems. It has recently been observed [36] that properly applying ADMM to difficult convex optimization problems results in fast algorithms able to exploit problem structure. Here, we apply ADMM to the PPR problem resulting in an algorithm which alternately solves *i) n* one-dimensional convex optimization problems and *ii)* the *ℓ*_2_ case of PPR. We start with the primal form (1) of PPR and replace the hard constraint **u** ≥ 0, **u**^*T*^𝟙_ ≤ 1 with the convex indicator function *g* where *g*(**u**) = 0 if **u** ≥ 0, **u** 𝟙≤ 1, and *g*(**u**) = ∞ otherwise. Then, we obtain the following equivalent form of the PPR problem:𝟙

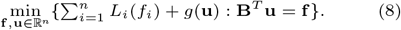

Following [35], we introduce the *augmented Lagrangian*

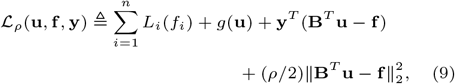

where *ρ >* 0 is a scalar hyperparameter and **y** *∈* ℝ^*n*^ is the dual variable. By Lagrangian duality theory, solving (8) is equivalent to finding a saddle point of ℒ_*ρ*_. To find such a point, ADMM employs an iterative algorithm which at each iteration, minimizes **f**, then **u**, and finally updates the dual variable **u** in an ascent step. Specifically, ADMM performs the following iterations starting at an initial solution (**u**^(0)^, **f** ^(0)^, **y**^(0)^):

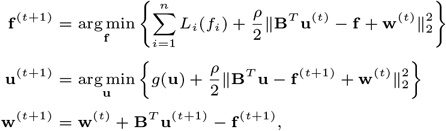

where 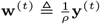 is the rescaled dual variable.

The **f**-update, by separability of the objective, consists in solving *n* one-dimensional convex minimization problems. The **u**-update consists of solving the PPR problem for the *ℓ*_2_ loss 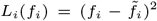 where 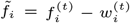 is a perturbation of the frequency at the *t*^th^ iteration. The **w**-update consists only of a matrix-vector product, which, by the combinatorial properties of **B** [22], takes 𝒪(*n*) time to perform.

##### A piecewise linear approximation approach

Approximation of non-linear functions with piecewise linear loss functions is a well-studied technique in both the optimization [37] and neural network literature [38], and is justified by a universal approximation theorem which states that arbitrary convex functions can be approximated by convex piecewise linear functions with a small number of pieces.

Given a set of differentiable, convex loss functions *L*_*i*_, we provide two algorithms for solving the PPR problem using piecewise linear approximation. The first algorithm, called *Piecewise Linear Approximation with k segments* (*PLA-k*), performs the following steps:

1. Select *k* − 1 breakpoints 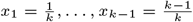.
2. Get 1^st^-order Taylor expansion *g*_*i*,*j*_ of *L*_*i*_ at each *x*_*j*_.
3. Define the piecewise linear approximation 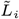 as the pointwise maximum over the Taylor approximations: 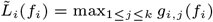.
4. Solve the PPR problem using TSDDP with piecewise linear loss functions 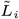.

From the definition of convexity, it follows that 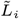 underestimates *L*_*i*_ and converges to *L*_*i*_ pointwise as *k* → ∞. The total runtime of PLA-*k* is dominated by the final step, leading to a time complexity of 𝒪(*nk* log^2^(*nk*)). Further, using a standard argument, we quantify the error of PLA-*k* for Lipschitz continuous loss functions *L*_*i*_, as stated in the following theorem.

###### Theorem 4.

*Let* 𝒯 *be a clonal tree on n mutations. Suppose L*_*i*_ *is β-Lipschitz continuous and* 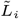 *was constructed using PLA-k. Then*, 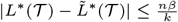

The second algorithm, called *Progressive Piecewise Linear Approximation* (*PPLA*), is slightly more sophisticated, instead *progressively* performing piecewise linear approximation, updating the frequency bounds at each step of the algorithm. Specifically, PPLA(*k, ρ, τ*) takes as input three parameters: (i) the number *k ∈* ℕ of segments, (ii) a multiplicative factor *ρ ∈* [0, 1] by which to adjust the search range, and (iii) a threshold parameter *τ ∈* [0, 1] used to terminate the procedure. The algorithm is as follows.

1. Set frequency bounds **f** ^−^ ≜ **0, f** ^+^ ≜ **1** and range *r* ≜ 1.
2. Obtain frequencies **f** by solving *k*-PLA with frequency lower bounds **f** ^−^ and upper bounds **f** ^+^.
3. Update range *r* ← *r · ρ*.
4. For each mutation *i ∈* [*n*]:
  i. Update frequency lower bound 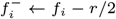.
  ii. Update frequency upper bound 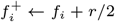.
5. If range *r > τ*, go to Step 2, otherwise return **f**.

Since the range *r* of PPLA(*k, ρ, τ*) shrinks by the constant factor *ρ* at every iteration of the algorithm, PPLA(*k, ρ, τ*) performs 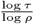 calls to *k*-PLA, yielding a total time complexity of 𝒪(*nk* log^2^ *nk ·* log *τ ·* log^−1^ *ρ*). Further, since PPLA(*k, ρ, τ*) improves the loss at every iteration, it enjoys the same convergence properties as *k*-PLA (Theorem 4). Practically, however, we have observed that PPLA(*k, ρ, τ*) finds more accurate solutions than *k*-PLA, albeit at the cost of increased runtime.

### fastppm algorithm

*fastppm* is an open-source implementation of the TSDDP algorithms for the *ℓ*_2_, binomial (via PLA, PPLA and ADDM) and beta-binomial (via PLA and PPLA) loss functions. *fastppm* is implemented in C**++** as both a shared library and command-line interface. *fastppm* provides Python bindings using PyBind. *fastppm* is available at: github.com/elkebir-group/fastppm.git.

## Results

### Fast and accurate regression using fastppm

We evaluated the performance of *fastppm* against five existing methods for the Perfect Phylogeny Regression problem under an *ℓ*_2_ and binomial loss. For both losses, we compared *fastppm* against the state-of-the-art convex programming solvers CVXOPT [39], ECOS [40], Mosek [41], and Clarabel [42]. For the *ℓ*_2_ loss, we additionally compared against the projectppm solver developed in [25, 29]. To evaluate each method, we simulated 90 tumor phylogenies, measuring the runtime and loss of the inferred frequencies (see Supplementary Methods for details). In addition to the observed frequencies (for *ℓ*_2_ loss) or the variant and total read counts (for binomial loss), all methods were provided the ground-truth simulated tree and were run in single-threaded mode on a 2.4 GHz CPU with 4 GB RAM.

In terms of the *ℓ*_2_ loss, *fastppm* was on average 111.1*×* faster than projectppm, ranging from 33.3*×* faster when the number of mutations was small (*n* = 100) to 163.2*×* faster when the number of mutations was large (*n* = 4000). Compared to general-purpose convex programming solvers, *fastppm* yielded another order of magnitude improvement, outperforming the commercial solver Mosek by 858.49*×* on average. Since the *ℓ*_2_ loss is strictly convex and all solvers exactly solve the optimization problem, the inferred objective values were identical.

For the binomial loss, conic programming solvers ECOS [40], Mosek [41], and Clarabel [42] struggled to terminate without error on the vast majority of instances (54/270, see Fig. S1). Although CVXOPT [39] terminated on all instances, all three modes of *fastppm* (Section 3.4) were several orders of magnitude faster. Specifically, *fastppm*-ADDM, *fastppm*-50-PLA, *fastppm*-100-PLA, and *fastppm*-PPLA were a mean of 400.1*×*, 43.8*×*, 22.3*×*, and 13.6*×* faster than CVXOPT, respectively. The loss of the solutions inferred by *fastppm*-ADMM, *fastppm*-50-PLA, and *fastppm*-100-PLA was at most 4% higher than that inferred by CVXOPT, while the loss inferred by *fastppm*-PPLA was similar or slightly lower than that inferred by CVXOPT (Fig. S2).

In summary, *fastppm* was at least an order of magnitude faster than existing methods while achieving nearly identical solutions on all problem instances. In addition, we compared the binomial and beta-binomial loss implemented using fastppm-PPLA on overdispersed simulation data, finding that using a beta-binomial loss better infers ground-truth frequencies (Fig. S3).

### Improved tree inference using **fastppm**

While the previous experiments assessed *fastppm* on a fixed ground-truth tree, we next evaluated *fastppm*’s ability to improve upon three existing phylogenetic inference pipelines: Sapling [26], CITUP [11] and Orchard [14]. Specifically, we modified the existing software for all three tools by replacing calls to existing solvers with calls to *fastppm*, obtaining three new tools: Sapling^*^, CITUP^*^ and Orchard^*^. The modifications consisted of fewer than fifty lines of code in each case. For each altered method, we compared the runtimes and distances to the ground-truth tree using two distance measures, the parent-child distance and the ancestor-descendant distance [43] (details in Supplementary Methods).

We obtained the strongest improvement when replacing the CVXOPT solver in Sapling [26] with *fastppm*-ADMM, where we obtained an average 406*×* improvement in runtime for *n* = 50 mutations and an increase in method accuracy (Fig. 3 and S4). Further, while Sapling was unable to infer phylogenies for instances beyond *n* = 50 mutations within a twelve hour time-limit, Sapling^*^ successfully inferred phylogenies on all instances with *n* = 500 mutations in under twelve hours. The next largest improvement was found with CITUP^*^, which obtained a 5-10*×* improvement in runtime and an improved rate of successful termination within a twelve-hour time limit over CITUP while achieving similar accuracy in recovering the true phylogeny (Fig. 3 and S5). Finally, Orchard^*^ obtained a 3-5*×* improvement in runtime while maintaining similar accuracy in recovering the true phylogeny as compared to Orchard (Fig. 3 and S6).

**Fig. 2.**
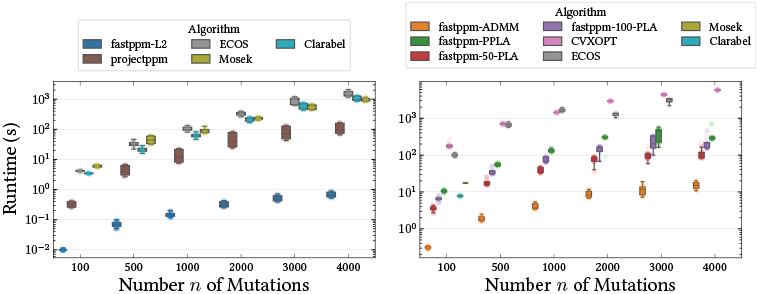
Wall-clock runtime of existing algorithms for the Perfect Phylogeny Regression problem across 90 tumor phylogenies for the **(left)** *ℓ*_2_ and **(right)** binomial loss functions. For the *ℓ*_2_ loss, all methods inferred the same optimal solution whereas Fig. S2 shows the objective value differences across methods for the binomial loss.

**Fig. 3.**
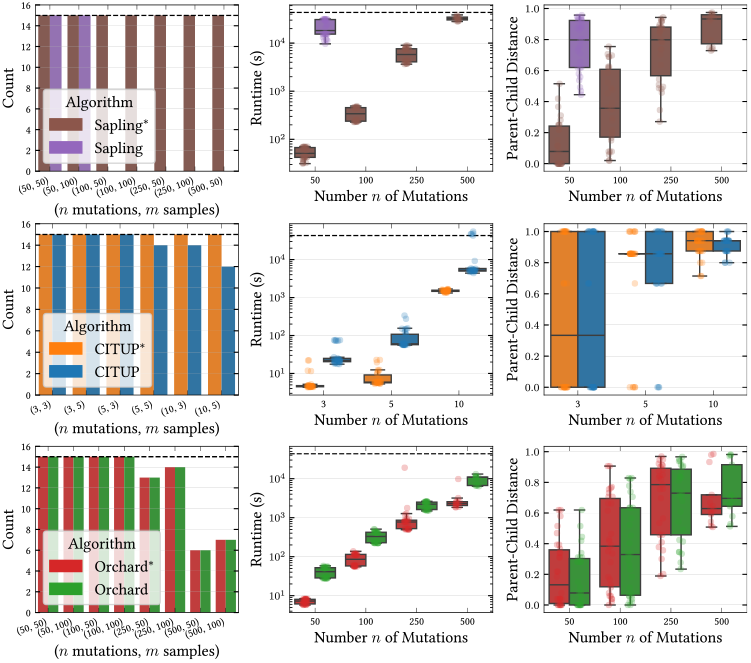
Number of successes within a twelve-hour time limit, wall-clock runtime (seconds), and tree inference error across three different methods. **(top)** Sapling^*^ is the result of replacing the CVXOPT solver for the binomial loss in Sapling [26] with *fastppm*-ADMM. **(middle)** CITUP^*^ is the result of replacing the CPLEX solver in CITUP [11] for the *ℓ*_2_ loss with *fastppm*-L2. **(bottom)** Orchard^*^ is the result of replacing the projectppm [25] solver in Orchard [14] for the *ℓ*_2_ loss with *fastppm*-L2.

In summary, we find that incorporating *fastppm* in existing tumor phylogeny inference methods provides substantial improvements in runtime, supporting larger instances with little to no degradation in performance across all three methods.

### Accurate phylogenetic reconstruction from low-coverage DNA sequencing data

To demonstrate the utility of directly modeling binomial loss rather than approximating the loss, we simulated 40 tumor phylogenies at 20*×* coverage and assessed how well the two top-performing methods, Orchard and Sapling^*^, recovered the true phylogeny. We hypothesized that because Orchard relies on the *ℓ*_2_ loss for phylogenetic inference, while Sapling^*^ directly models read counts via the binomial loss, the latter would produce more accurate phylogenies on such shallow sequencing data.

Confirming our hypothesis, we found that Sapling^*^ outperformed Orchard in terms of both runtime and reconstruction accuracy on low coverage simulated bulk DNA sequencing data (Fig. 4). For example, on low-coverage data with *n* = 50 mutations and *m* = 50 samples, Orchard inferred phylogenies with a mean (normalized) ancestor-descendant (AD) distance of 0.21, while the phylogenies inferred by Sapling^*^ had a mean AD distance of 0.09. As the number of mutations increased, the trend held, with the performance of both methods degrading as the number of mutations increased (Fig. 4). In terms of runtime, Sapling^*^ was at least twice as fast as Orchard across all instances. However, without the improvement in runtime due to *fastppm*-ADMM, the runtime of Sapling^*^ would have been intractable for most use cases. In particular, processing instances with only *n* = 250 mutations and *m* = 50 samples would have taken an estimated 38 hours using CVXOPT, in contrast to an average of 10 minutes with *fastppm*-ADMM.

**Fig. 4.**
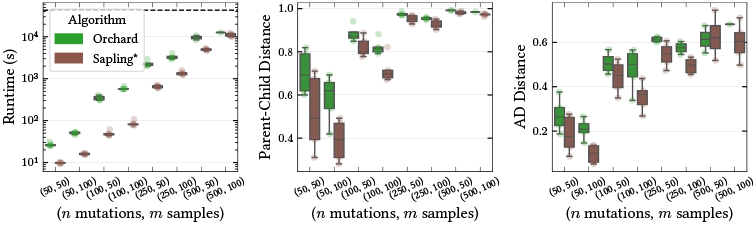
Wall-clock runtime (s), parent-child distance, and ancestor-descendant distance for the phylogenies inferred by Orchard [14] and Sapling^*^ on simulated 20*×* coverage bulk DNA sequencing data.

### Improved phylogenetic inference on a mouse model of colorectal cancer

Finally, we analyzed the patient-derived xenograft POP66 from a model of colorectal cancer in mouse, from which *m* = 8 bulk samples underwent whole-exome sequencing at 50*×* coverage [44]. Following the analysis in the original publication, we excluded mutations that were contained in copy number impacted regions and used the copy number corrected read counts provided by [44]. After copy number correction, *n* = 65 mutations were used for the analysis in [44]. To test the effect of directly modeling the binomial loss, rather than using the variant and total read counts directly we down-sampled the total read counts to 20*×* coverage (Supplementary Methods). Then, we applied Orchard, which uses the *ℓ*_2_ loss, and Sapling^*^, which uses *fastppm*-ADMM for the binomial loss, to the down-sampled data and compared the inferred frequencies to those inferred on the full data.

Both methods completed tree inference in minutes, with Sapling^*^ taking 3 minutes and 5 seconds and Orchard taking 7 minutes and 51 seconds when parallelized across a 16 core CPU. Notably, the original Sapling algorithm (which uses CVXOPT rather than *fastppm*-ADMM) would have taken over 22 hours. The phylogeny inferred by Sapling^*^ better recapitulated the down-sampled and original data compared to the phylogeny inferred by Orchard (Table 2). In more detail, we computed the *ℓ*_2_ error and the binomial loss for the frequency matrices 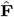 inferred by each method against the down-sampled and original data. For the *ℓ*_2_ error metric, the frequencies 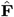 inferred by Sapling^*^ on the down-sampled data were 20% closer to the observed frequencies 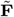 on the original data (Table 2). In terms of the binomial likelihood, the frequencies 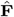 inferred by Sapling^*^ better explained (NLL: 10720.6) the observed variant reads as compared to those inferred by Orchard (NLL: 10801.9) (Table 2). Qualitatively, the phylogenies inferred by the two methods differed substantially (Parent-Child Distance: 0.84, Ancestor-Descendant Distance: 0.29). For example, while the Orchard [14] phylogeny had a truncal CNR1 mutation, the Sapling^*^ phylogeny had a truncal KDR mutation (Fig. S7-S8). However, the KDR and CNR1 mutations occurred relatively early in both phylogenies, suggesting a modest level of concordance.

**Table 2.**
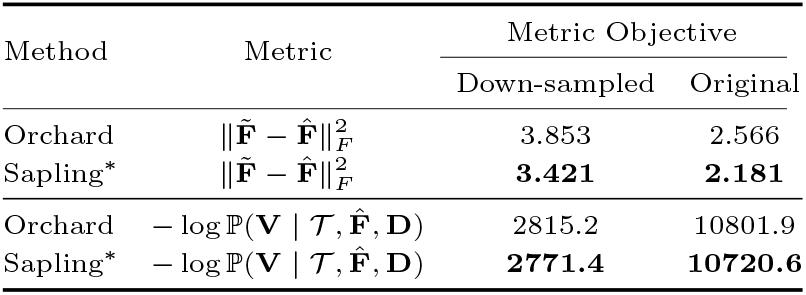
Analysis of POP66 patient derived xenograft data modeling colorectal cancer in mice [44]. The fit of the frequency matrices 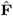 inferred by Orchard [14] and Sapling^*^ to the original (resp. down-sampled) variant read count matrix **V**, total read count matrix **D**, and observed frequency matrix 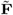.

These findings demonstrate that *fastppm* enables the application of more accurate loss functions, whose effect becomes especially prominent in shallow coverage settings.

## Discussion

We introduced TSDDP, a new algorithmic technique capable of solving the Perfect Phylogeny Regression problem for arbitrary convex loss functions. Using TSDDP, we built *fastppm*, an easy-to-use tool which supports several convex loss functions, including the *ℓ*_2_, piecewise linear, binomial and beta-binomial loss. As future work, the implementation *fastppm* does not leverage the TSDDP algorithm’s ability to efficiently recompute the optimal objective upon slight perturbations to the tree topology. Supporting re-optimization of the PPR problem would yield additional speedups of existing tumor phylogeny inference pipelines, which iteratively update candidate topologies via tree moves such as subtree prune and regraft (SPR) [13, 14, 26]. Second, while the PLA-*k* algorithm appears to construct an optimal piecewise linear approximation when the loss function is not-known beforehand, PLA-*k* could be improved by accounting for the structure of specific loss functions. Finally, we envision extending beyond the framework of maximum likelihood, instead computing the marginal likelihood with an integral over **f** respecting the (SC).

## Supplementary Results

We first state two elementary observations which hold for all loss functions *L*_*i*_ and will be used in the following sections.

### Observation 1.

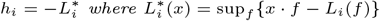 *is the convex conjugate of L*_*i*_.

### Observation 2.

*J*_*i*_ *is concave*.

### The case of quadratic loss

In the case of a weighted quadratic loss we have 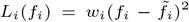, where 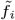 denotes the observed frequencies and *w*_*i*_ ≥ 0 denotes the weight assigned to the *i*^th^ frequency. Since 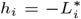, where 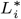 is the convex conjugate of the quadratic loss *L*_i_, we have 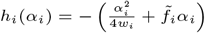.When *i* is a leaf, 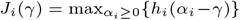,and by characterizing the maximizer in terms of the derivative, we have

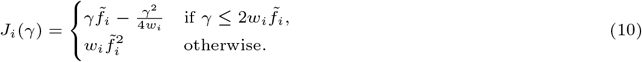

In this case, *J*_*i*_ has a piecewise linear derivative and as we will soon show, *J*_*i*_ always has a piecewise linear derivative.

Consequently, we fix (i) the computational representation *ℛ* (*J*_*i*_) to consist of the value *J*_*i*_(0) along with the slopes, intercepts, and breakpoints of the derivative 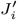.In this representation, the value *J*_*i*_(0) is stored as a scalar, while the slopes, breakpoints and intercepts of 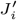 are stored in three one-dimensional arrays sorted in ascending order of the breakpoints. To compute the representation (ii) of the leaves, it suffices to use (10) to obtain the representation of *J*_*i*_, taking 𝒪(1) time since 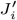 has only a single breakpoint.

It remains to show that *J*_*i*_ always has a piecewise linear derivative and describe how to compute the representation *ℛ* (*J*_*i*_) at an internal node *i*. To do this, we assume that the inductive hypothesis (i.e. *J*_*i*_ has a piecewise linear derivative) holds for all descendants of node *i*. By the base case given in (10), the inductive hypothesis holds for the deepest, non-leaf node, allowing us to propagate the claim up the tree.

Assuming that for all descendants *j* of node *i*, the representations *ℛ* (*J*_*j*_) are available and 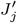 is a piecewise linear function, we first characterize the maximizer in *J*_*i*_(*γ*),

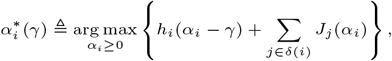

where we let arg max denote the unique, left most maximizer. Since the inner term is concave by (Observation 2), it follows that

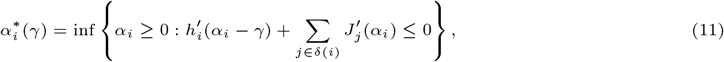

where we take the infimum of the empty set to be 0. The reason for studying the arg max rather than the max is that the derivative of a (piecewise) quadratic function is (piecewise) linear, and consequently easier to work with.

Next, assuming that the derivatives 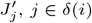, have at most *k*_*j*_ pieces with 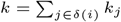, we write the sum

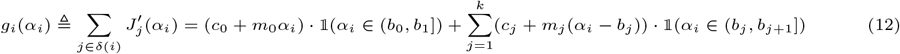

where −∞ = *b*_0_ *< b*_1_ *<* … *< b*_*k*_ *< b*_*k*+1_ = +∞ are the *k* distinct breakpoints of the piecewise linear functions 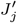.As 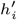 is affine, we have that

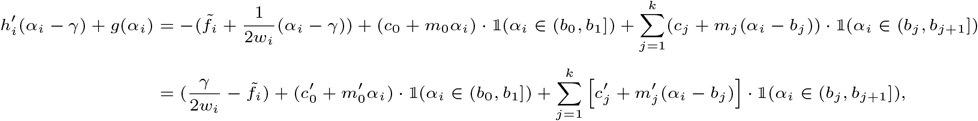

where 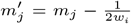 for 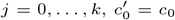,and 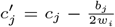 for *j* = 1, …, *k*. Then, 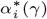 is the smallest point *α*_*i*_ ≥ 0 such that

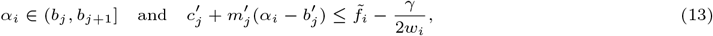

where 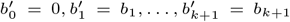.We can now use this relation to show that 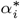 is linear on certain pieces, formalized as follows.

#### Lemma 1.

*If* 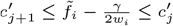, *then* 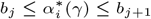.

*Proof* If *α*_*i*_ *< b*_*j*_, we have *α*_*i*_ *∈* (*b*_*l*_, *b*_*l*+1_] for some *l < j*. However, since 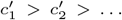 from the fact that *g*_*i*_(*α*) is decreasing (as *J*_*i*_ is concave), this implies that 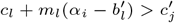.Consequently, *α*_*i*_ could not satisfy (13) if *α*_*i*_ *< b*_*j*_. In the other direction, since *α*_*i*_ = *b*_*j*+1_ satisfies (13), no *α*_*i*_ *> b*_*j*+1_ can be the smallest such *α*_*i*_ satisfying (13). This completes the proof. □

Since 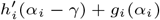 is linear on each piece [*b*_*j*_, *b*_*j*+1_], we derive the following explicit form of 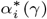,completing our characterization of 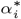.

#### Lemma 2.

*Let* 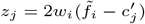.*Then*,

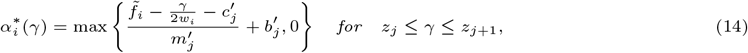

*Proof* Suppose *z*_*j*_ ≤ *γ* ≤ *z*_*j*+1_. Then, if 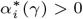,by Lemma 1 we have that

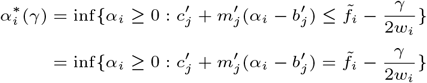

where the first equality follows from the assumption that *z*_*j*_ ≤ *γ* ≤ *z*_*j*+1_ and the second from the linearity of *h*_*i*_(*α*_*i*_ − *γ*) + *g*_*i*_(*α*_*i*_) when *α*_*i*_ *∈* [*b*_*j*_, *b*_*j*+1_]. Solving the equality for 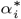 and taking care of non-negativity completes the proof. □

Now, we are ready to obtain an explicit form for 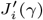. In particular, we have the following form for 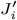.

#### Lemma 3.

*Let* 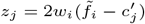 *and suppose γ ∈* [*z*_*j*_, *z*_*j*+1_] *with b*_*j*_ ≥ 0. *Then*,

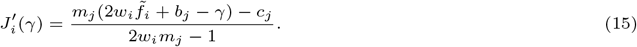

*Further*, 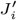 *is piecewise linear*.

*Proof* Using the chain rule for subgradients, we have that

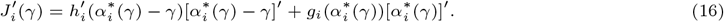

Since 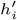 is affine, an explicit expression for the first term is straightforward to compute. Next, we note that by Lemma 1, 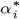 is piecewise linear. Since *g*_*i*_ is also piecewise linear, this implies that 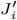 is piecewise linear with a finite number of pieces.

Next, note that 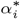 is monotonically increasing: to see this, observe 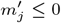 by the concavity of *J*_*i*_ and 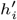,and use the expression given in Lemma 1. A quick computation then reveals 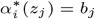,provided *b*_*j*_ ≥ 0, implying 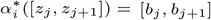.Using (16), it then follows that 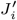 is also linear when *γ ∈* [*z*_*j*_, *z*_*j*+1_], since then *α*_*i*_(*γ*) *∈* [*b*_*j*_, *b*_*j*+1_]. Taking advantage of this linearity, plugging in (14) to (16), and performing algebraic manipulations verified using the computer algebra system Mathematica v14.1, we obtain (15). □

Setting *l* = min{*j* : *b*_*j*_ ≥ 0}, we have obtained an explicit form for 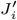 for all *γ* ≥ *z*_*l*_. In particular, from *z*_*l*_ to 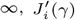 is piecewise linear with breakpoints *z*_*l*_ ≤ … ≤ *z*_*k*_ and slopes and intercepts given by (15). Thus, in 𝒪(*k*) time we are able to compute the slopes, intercepts and breakpoints of 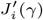 for *γ* ≥ *z*_*l*_, nearly completing (iii) of the TDSPP algorithm. To the left of *z*_*l*_, there can be at most one additional breakpoint, which is somewhere on the interval [*b*_*l*−1_, *b*_*l*_]. By a similar line of reasoning, we can find this breakpoint, called *x*, using Lemma 1. Clearly, for 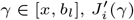 is again linear, with its slope and intercept easily computed. Since for all 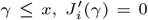, we have completely described the slopes and intercepts of 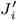 using only 𝒪(*k*) time. Further, we have shown that the total number of distinct breakpoints in 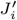 is bounded by *k*, the total number of breakpoints in the functions 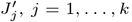.We are now ready to complete the proof of Theorem 2.

*Proof of Theorem 2*. For the base case (ii) of a tree with a single node it suffices to compute the representation of a single leaf, which by (10) takes 𝒪(1) time.

For the inductive step (iii), consider a tree 𝒯 with *n* nodes and *l* = |*δ*(*r*)| subtrees 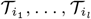 rooted at the children *δ*(*r*) = {*i*_1_, …, *i*_*l*_} of the root vertex *r* of *T* and suppose the representations 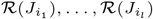 have already been computed. To compute *ℛ* (*J*_*i*_), we first compute 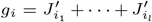, which consists of the summation of *l* piecewise linear functions with a total of *n* − 1 breakpoints, following the preceding arguments. This summation can be performed in at most *C*_1_*n* log(*l*) time for some constant *C*_1_ by recursively merging the *l* arrays of sorted breakpoints in the representations 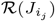 while keeping track of the associated slopes. Then, using Lemma 3 and the preceding arguments, we compute 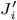 using an additional *C*_2_*n* time for some constant *C*_2_. To compute the intercept *J*_*i*_(0) also takes *C*_2_*n* time, using the equality *J*_*i*_(0) = 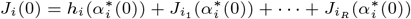.

Adding the time taken at each node, it takes a total of *T* time to compute the representation ℛ(*J*_*r*_) of the root vertex:

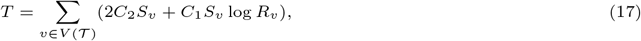

where *S*_*v*_ = |*V* (𝒯_*v*_)| is the size of the subtree rooted at *v* and *R*_*v*_ = |*δ*(*v*)| is the number of children of vertex *v*. We bound *T* using Holder’s inequality in two different ways. First, we have

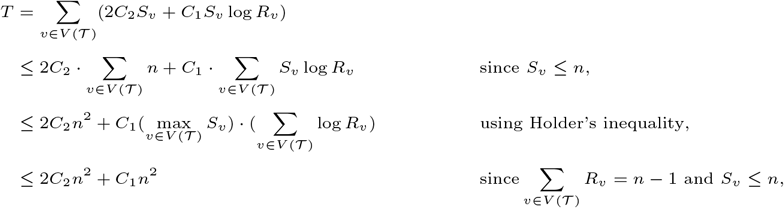

yielding a runtime bound of 𝒪(*n*^2^). Second, let 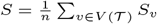, we then have

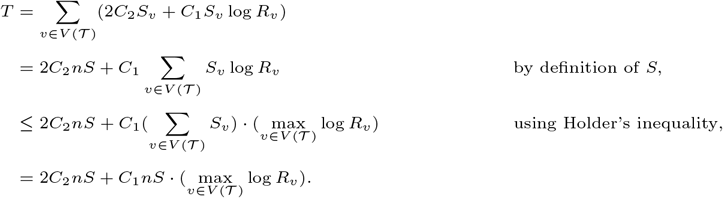

This yields the second runtime bound of 𝒪(*nS* log *R*) = 𝒪(*Sn* log *R*). However, it is well known that 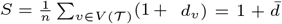 where *d*_*v*_ is the distance of *v* from the root to *v*. This is because in computing 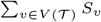, each node *v* is counted a total of *d*_*v*_ + 1 times, once for each ancestor of *v*. Consequently, we obtain a third bound of 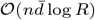,completing the proof.

The case of convex piecewise linear loss

In the case of a convex, piecewise linear loss function with *k* breakpoints, we have

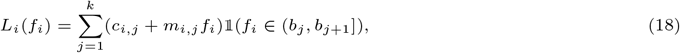

where −∞ = *b*_0_ *< b*_1_ *< b*_2_ *<* … *< b*_*k*+1_ = +∞ are the *k* distinct breakpoints of *L*_*i*_. To design our TSDDP algorithm for this class of loss functions, we show that steps (i)-(iv) consist solely of the operations of addition, conjugation, and infimal convolution [31] of convex piecewise linear functions, for which specialized algorithms and data structures have been developed [34].

Specifically, given two convex functions *f, g* : ℝ → ℝ ∪ {*±*∞}, we denote by *f* + *g* the sum, by *f* □ *g* the *infimal convolution*, and by *f*^*^ the *convex conjugate* which are defined pointwise as,

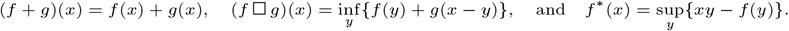

Importantly, we note that the infimal convolution is related to conjugation through the identity

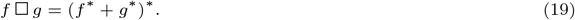

Under this notation, it follows that both steps (ii) and (iii) of the TSDDP algorithm can be written in terms of the three aforementioned operations. Namely, for a vertex *i*, we have that

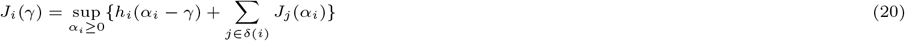

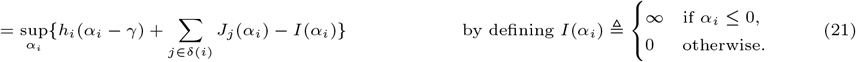

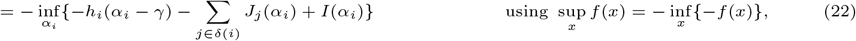

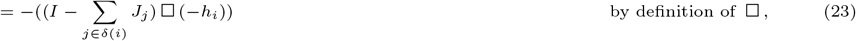

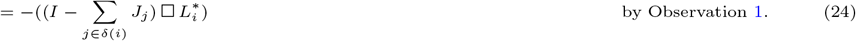

That is, computing the representation *ℛ*(*J*_*i*_) is equivalent to computing the representation after performing the operations of summation and conjugation, followed by infimal convolution.

In more detail, at a leaf vertex *i*, it follows from (24) that 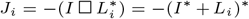, which is the negation of the conjugation of a convex, piecewise linear function. From Observation 1 in [34], it follows *J*_*i*_ is concave and piecewise linear with *k* breakpoints, where the breakpoints of *L*_*i*_ become the slopes of *J*_*i*_ and the slopes of *L*_*i*_ become the breakpoints of *J*. Inductively assuming that *J*_j_ are concave and piecewise linear for all children *j ∈ δ*(*i*), we have that 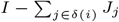 is convex and piecewise linear. Again applying Observation 1 from [34], it follows that 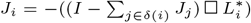 is concave and piecewise linear.

Since *J*_*i*_ is a concave, piecewise linear function, we fix (i) the representation |*J*_*i*_) to consist of the slopes, breakpoints, and intercept of *J*_*i*_. However, rather than näively storing the slopes and breakpoints in an array or a linked list, we use the representation described in [34] which represents the breakpoints and slopes in a balanced binary search tree by storing the *difference* in the slopes and breakpoints on the nodes of the tree. Indeed, using such a representation, [34] proves (see Proposition 6.1) that obtaining the representation of the sum of *l* concave, piecewise linear functions with *k*_1_ ≤ … ≤ *k*_*l*_ breakpoints, 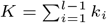,takes 𝒪(*K* log *K*) time. Similarly, [34] proves that obtaining the representation of the infimal convolution of two piecewise linear functions with *K* and *K*^*′*^ breakpoints takes 𝒪(min{*K, K*^*′*^} log^2^(*K* + *K*^*′*^)) time.

We are now ready to prove Theorem 3.

*Proof of Theorem 3*. Suppose inductively that for all trees 𝒯 of size less than *n*^*′*^ *< n* we can compute the representation ℛ (*J*_|*T*)_) in 𝒪(*n*^*′*^ log^2^(*n*^*′*^*k*)) time. This holds for the base case (ii) of a tree with a single node, since it then suffices to compute the representation of a single leaf, which by the preceding arguments takes 𝒪(*k* log (*k*)) time.

Next, consider a tree 𝒯 with *n* nodes and *R* = |*δ*(*r*)| subtrees 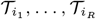 rooted at the children *δ*(*r*) = {*i*_1_, …, *i*_*R*_} of the root vertex *r* of 𝒯. By the inductive hypothesis, we can compute the representations 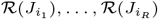 in 𝒪((*n* − 1)*k* log (*nk*)) time. By the preceding arguments, obtaining the representation *ℛ* (*J*_*r*_) consists of first obtaining the representation of 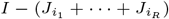, which takes 𝒪((*n* − 1)*k* log (*nk*)) time as the total number of breakpoints in 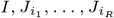 is bounded by (*n*−1)*k*+1. Computing the representation of the convolution 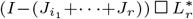 requires only an additional 𝒪(*k* log^2^(*nk*) time as *L*^*^ only has *k* pieces. Consequently, the time to compute the representation *ℛ* (*J*_*r*_) is 𝒪(*nk* log^2^(*nk*)) in total.

Observing that a concave, piecewise linear function must be maximized at a breakpoint, we solve the one-dimensional optimization problem (iv) in 𝒪(*nk*) time by scanning over the breakpoints. This completes the proof. □

### Convexity of the negative beta-binomial log-likelihood

The beta-binomial distribution, BetaBin(*d, α, β*), is a compound distribution described in two steps. First, the binomial proportion *p* is drawn from a beta distribution parameterized by shape parameters *α, β*, i.e. *p* ~ Beta(*α, β*). In our setting, the mean of this beta distribution corresponds to the frequency *f*, i.e. *f* = *α/*(*α* + *β*). Second, the number of successes, or in our case the number of variant reads *v*, are drawn from a binomial distribution with *d* trials and success probability *p*, i.e. *v* ~ Bin(*d, p*). The probability mass function then equals:

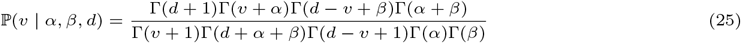

where Γ(*x*) is the gamma function.

We use a re-parametrization where, rather than the two shape parameters *α, β*, we are given a precision parameter *s* = *α* + *β*. In our setting, in addition to *s*, we are given the total number *d* of reads (or trials) and the variant number *v* of reads (or successes). The frequency *f* (or mean of the beta distribution) is the parameter of interest in our setting.

Since *s* = *α* + *β* and *f* = *α/*(*α* + *β*) = *α/s*, we have *α* = *fs* and *β* = *s* − *fs*, which allows us to rewrite (25) as

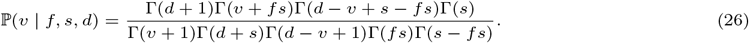

Ignoring constant terms with respect to the parameter *f*, the negative log-likelihood is up to an additive constant-factor then equal to:

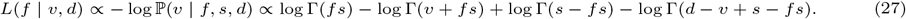

Convexity of *L*(*f* | *v, d*) with respect to *f* can be shown using the monotonicity (decreasing) of the trigamma function (i.e. 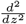 ln Γ(*z*)).

## Supplementary Methods

### Simulation details

Each simulation was defined by a tuple of four parameters (*n, m, c, r*), as described below.

*n*: the number of mutations/nodes in 𝒯.

*m*: the number of samples.

*c*: the expected number of total reads or *coverage*.

*r*: the random seed.

The output of each simulation is a clonal tree 𝒯 on *n* mutations, a *n*-by-*n* clonal matrix **B**, a *m*-by-*n* variant read count matrix **V**, a *m*-by-*n*-total read count matrix **D**, a *m*-by-*n* frequency matrix **F**, and an observed frequency matrix 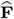.

To construct a simulated instance for a fixed set of parameters (*n, m, c, r*), we first uniformly at random sampled, using Wilson’s algorithm [45], a clonal tree 𝒯 with *n* mutations and constructed the associated clonal matrix **B**. Then, for each of the *m* samples, we sampled a usage vector 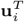 from a Dirichlet distribution, stacked them into a usage matrix **U** = [**u**_*i*_], and fixed the frequency matrix as **F** = **UB**. To generate the observed read counts, we sampled the total number *d*_*ij*_ of reads for the *j*^th^ mutation in the *i*^th^ sample using *d*_*ij*_ ~ Poisson(*c*). Then, using the total read counts, the variant reads were sampled using *v*_*ij*_ ~ Binomial(*d*_*ij*_, *f*_*ij*_). To obtain variant and total read count matrices, we set **V** = [*v*_*ij*_] and **D** = [*d*_*ij*_]. The observed frequency matrix was set to 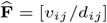.

### Executing and evaluating regression algorithms

We evaluated five existing regression algorithms for the *ℓ*_2_ and binomial negative log-likelihood loss against *fastppm* on 90 simulated tumor phylogenies containing *n ∈* {100, 500, 1000, 2000, 3000, 4000} mutations and a read coverage of *c ∈* {30, 100, 1000} across *r ∈* {1, 2, 3, 4, 5} random number generator seeds. For the *ℓ*_2_ loss, we evaluated the specialized regression algorithm *projectppm* [25]. In addition, we evaluated state-of-the-art convex optimization solvers CVXOPT [39], ECOS [40], Mosek [41], and Clarabel [42] on both the *ℓ*_2_ and binomial log-likehood loss.

To build *projectppm* [25], we compiled the software using GCC with optimization flags -O3 and `-ffast-math`. Then, to execute the software, we passed in the observed frequency matrix 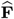 and the ground truth clonal tree *T* to the solver. Additionally, we specified that the solver only output the optimal objective value and not the inferred usage and frequency matrices. Finally, to time the software, we used the Linux tool `/usr/bin/time`, which provided a sufficiently high precision clock to benchmark the solver runtime.

To evaluate conic optimization solvers ECOS [40], Mosek [41], and Clarabel [42], we accessed the solvers through the Python convex optimization interface CVXPY [46]. As the objective was trivially separable across the *m* samples, to ensure a fair comparison, we passed each sample one at a time using CVXPY’s *Parameter* object. Then, for each sample *p*, we setup the following optimization problem in CVXPY:

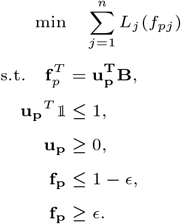

For the negative binomial log-likelihood loss, we fixed *ϵ* = 10^−5^ to avoid numerical stability issues which persisted across all solvers, while we set *ϵ* = 0 for the *ℓ*_2_ loss. For the negative binomial log-likelihood loss, we set 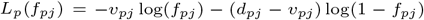. For the *ℓ*_2_ loss, we set 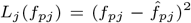.To benchmark the solvers, we recorded the solve time as provided by the solver and ignored the time spent building the model and the time spent in CVXPY’s interface. Since each sample was processed independently, we summed the runtime and objective across all *m* samples to obtain the final runtime and objective.

To evaluate convex optimization solver CVXOPT [39], we used the non-linear objective convex optimization solver `cvxopt.solvers.cp` provided through the Python interface. To use the solver, we passed in a helper function which computes the objective *L*(**F**), the gradient ∇*L*(**F**), and the Hessian matrix ∇^2^ *L*(**F**) of the objective

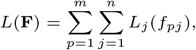

with respect to the flattened frequency matrix vec **F**. Since the Hessian matrix is diagonal, we returned a sparse representation of the *nm*-by-*nm* Hessian matrix. In total, computing the objective, gradient, and Hessian matrix took 𝒪(*nm*) time. To benchmark the solver, we used the runtime and objective value reported by the solver.

All solvers were ran with default parameters, excepting an increased number of maximum iterations for the Clarabel and ECOS solvers to improve the solvers’ success rate. An execution of a solver was called successful if for all *m* samples, the solver successfully terminated and emitted a finite objective value.

### Executing and evaluating tree inference algorithms

To evaluate the resulting six methods, we simulated tumor phylogenies using an identical procedure as in Simulation details, but varied the parameter settings to fit each method. For Sapling and Orchard, which progressively grow clonal trees, we simulated 120 tumor phylogenies with *n ∈* {50, 100, 250, 500} mutations, *m ∈* {50, 100} samples, and *c ∈* {30, 100, 1000} read coverage across *s ∈* {1, …, 5} random number generator seeds. For CITUP, which exhaustively enumerates all rooted, unlabeled trees with *n* nodes, we simulated 180 tumor phylogenies with *n ∈* {3, 5, 10} mutations, *m ∈* {3, 5} samples, and *c ∈* {30, 100, 1000} read coverage across *s ∈* {1, …, 5} random number generator seeds. When running the six methods, we provided Sapling and Sapling^*^ with a single core and four GB of memory, Orchard and Orchard^*^ with eight cores, and four GB of memory and CITUP and CITUP^*^ with sixteen cores and four GB of memory on 2.4 GhZ CPUs.

For each method and simulated instance, we measured the wall-clock runtime and the accuracy in recovering the ground truth tumor phylogeny, quantified in terms of both the normalized parent-child distance and the normalized ancestor-descendant distance. The (normalized) parent-child distance [26] between two directed graphs *T*_1_, *T*_2_ on the same vertex set *V* (𝒯_1_) = *V* (𝒯_2_) is the normalized symmetric difference of their edge sets. Namely,

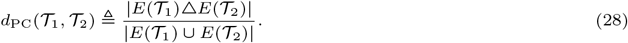

The (normalized) ancestor-descendant distance [43] is then defined as the parent-child distance between the transitive closures cl(*T*_1_), cl(*T*_2_). Namely,

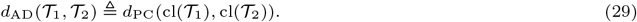

Both metrics take values in the interval [0, 1], with *d*(𝒯_1_, 𝒯_2_) = 0 if and only if 𝒯_1_ = 𝒯_2_.

### Executing and evaluating Orchard and Sapling^*^ on low-coverage DNA sequencing data

To evaluate Orchard and Sapling^*^, we simulated tumor phylogenies using an identical procedure as in Simulation details, but modified the simulation parameters and the tree moves in Sapling^*^. Specifically, we simulated 40 tumor phylogenies with *n ∈* {50, 100, 250, 500} mutations, *m ∈* {50, 100} samples, and *c* = 20 read coverage across *s ∈* {1, …, 5} random number generator seeds. To modify the tree moves in Sapling^*^, we considered the expanded set of moves used in Orchard [14] and fastBE [22], which upon the addition of a new mutation to the current tree, considered all possible (in worst-case 2^*n*^) insertions that respected the current tree’s partial order.

### Down-sampling read counts on a mouse model of colorectal cancer

To down-sample the POP66 colorectal data to a lower 20*×* coverage, we took the original variant and total read count matrices **V, D** and constructed the observed frequency matrix 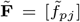 where 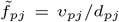.Then, we generated a synthetic *m*-by-*n* total read count matrix 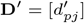 by drawing 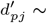 Poisson(20). Using the total read count matrix, we then constructed the down-sampled variant matrix 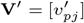 by drawing the variant reads using the observed frequencies: 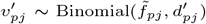. The result of this process was the original variant and total read count matrices **V, D** along with the down-sampled variant and total read count matrices **V**^*′*^, **D**^*′*^.

For the subsequent analysis, we built phylogenies by passing in the down-sampled read count matrices **V**^*′*^ and **D**^*′*^ to Orchard and Sapling^*^, obtaining tumor phylogenies 𝒯_1_ and 𝒯_2_ respectively. We then measured the concordance of both trees with the original and down-sampled data by computing *L*^*^(𝒯_*i*_ | **V, D**) and *L*^*^(𝒯_*i*_ | **V**^*′*^, **D**^*′*^) for *i* = 1, 2 under both the negative binomial log-likelihood and *ℓ*_2_ loss (Table 2).

## Supplementary Proofs

*Proof of Eq*. (6). The recurrence relation in (6) follows from the following chain of equalities:

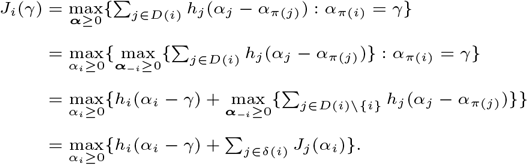

The first equality holds by definition of *J*_*i*_(*·*), the second equality holds by defining ***α***_−*i*_ = (*α*_0_, …, *α*_*i*−1_, *α*_*i*+1_, …, *α*_*n*_), the third equality holds by constancy of *h*_*i*_(*·*), and the final equality holds by the separability of the sum with *α*_*i*_ fixed. □

*Proof of Observation 2*. This follows from the stronger statement that *f* (*x*) = max_*y∈C*_ {*f*_1_(*y* − *x*) + *f*_2_(*y*)} is concave, provided that *f*_1_ and *f*_2_ are concave, and *C* is convex. To see this, observe that *f* (*x*) is the suprememal projection of the concave function *f*_1_(*y* − *x*) + *f*_2_(*y*) onto the *y* coordinate, over a convex set. It is well-known that supremal projections of concave functions (equivalently infimal projections for convex functions) preserve concavity. See equation (3.16) in Boyd [31] for a reference and proof.

Using the preceding result, the observation follows from induction over 𝒯. In the case where *i* is a leaf, 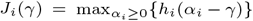 and the result holds by setting *f*_1_ = *h*_*i*_ and *f*_2_ = 0. Since concavity is preserved under summation, inductively, we have 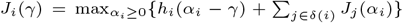 is concave upon setting *f* = *h* and 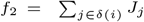.□

*Proof of Theorem 4*. Let 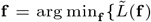 : **f** satisfies (SC)} and **f** ^*′*^ = arg min_**f**_ {*L*(**f**) : **f** satisfies (SC)}. Let **f** ^*′′*^ be the nearest point (under the ∥*·*∥_1_ norm) to **f** on the *n*-dimensional grid 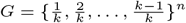.By construction, 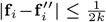.

Consequently,

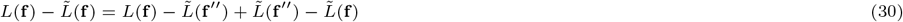

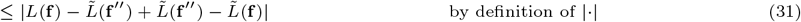

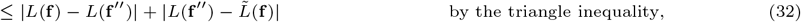

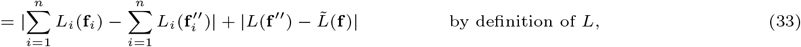

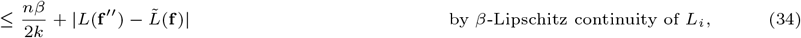

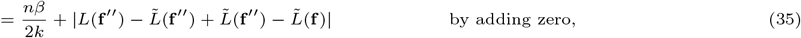

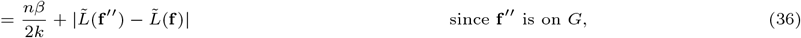

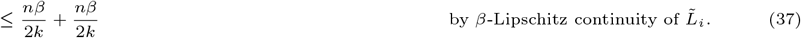

Using both the fact that 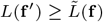 and *L*(**f** ^*′*^) ≤ *L*(**f**), we have

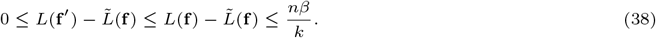

Consequently, 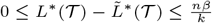, completing the proof. □

## Supplementary Figures

**Fig. S1.**
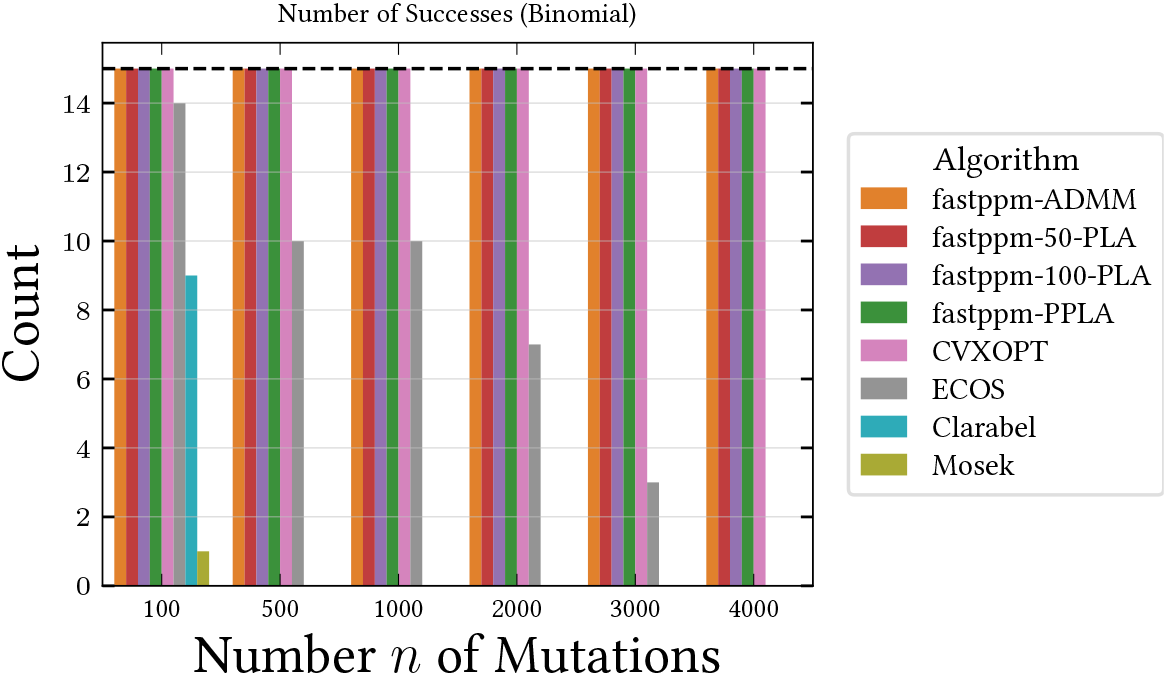
Number of successes of existing algorithms for the Perfect Phylogeny Regression problem across 90 tumor phylogenies for the binomial negative log-likelihood loss function. For the *ℓ*_2_ loss, all algorithms terminated successfully across all 90 instances and are excluded.

**Fig. S2.**
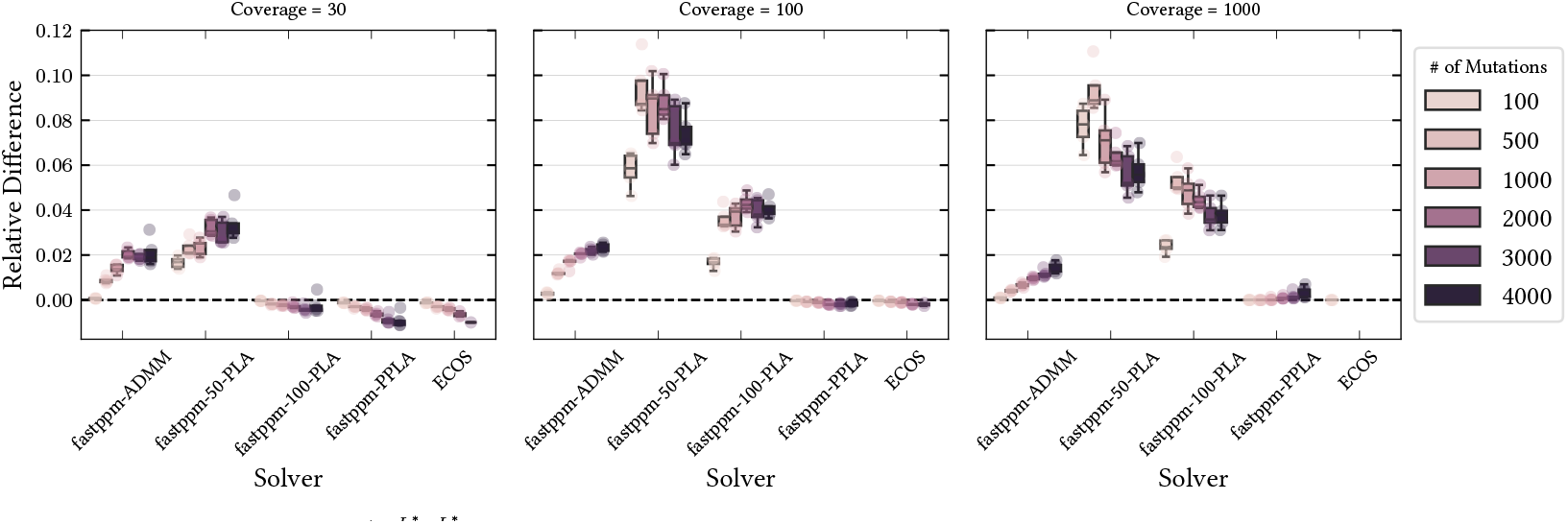
Relative difference 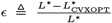 in the negative binomial log-likelihood of the inferred frequencies between all solvers and CVXOPT. *L*^*^ denotes the negative binomial log-likelihood of the inferred frequencies for a particular solver and 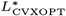 denotes the negative binomial log-likelihood of the inferred frequencies for CVXOPT.

**Fig. S3.**
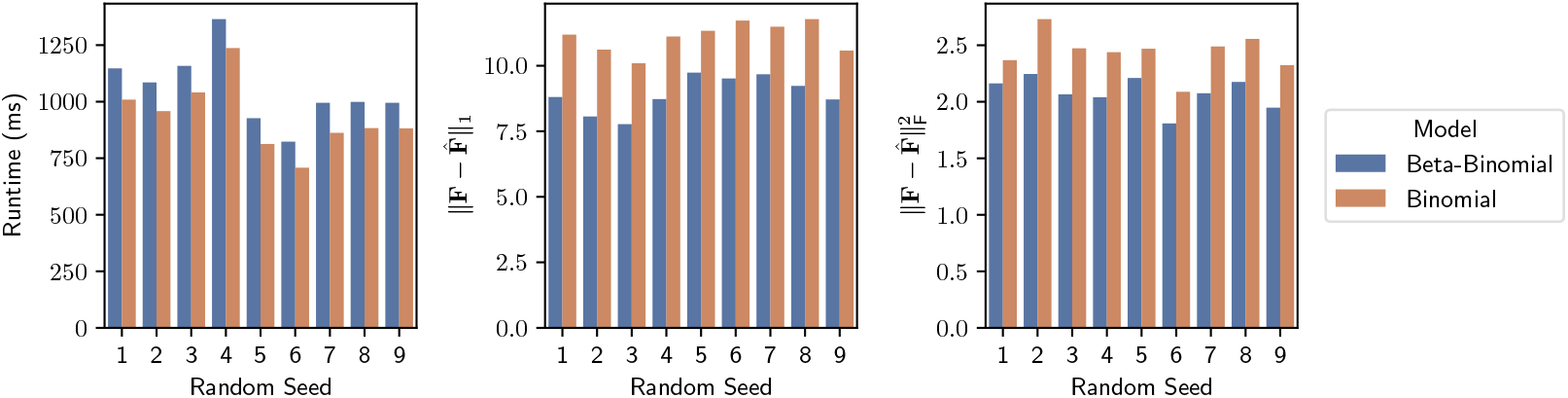
Runtime (ms) of PPLA and difference in the inferred and true frequency matrices when inferring frequencies using a beta-binomial and binomial loss function. Simulated data has *n* = 50 mutations, *s* = 100 samples, and *c* = 20 coverage. Variant read counts were generated using the beta-binomial distribution with precision parameter *s* = 2.0.

**Fig. S4.**
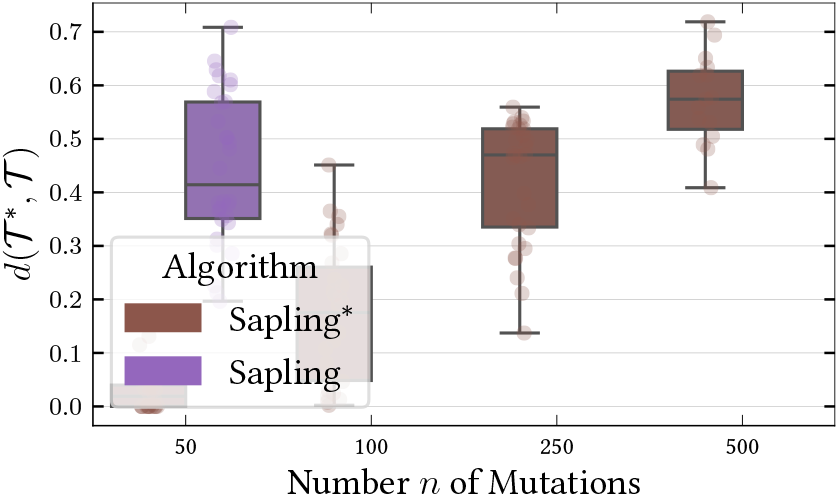
Ancestor-descendant distance between true and inferred phylogenies of Sapling [26] and Sapling^*^ across 120 simulated tumor phylogenies. Sapling^*^ is the result of replacing the CVXOPT solver for the binomial negative log-likelihood loss in Sapling with *fastppm*-ADMM.

**Fig. S5.**
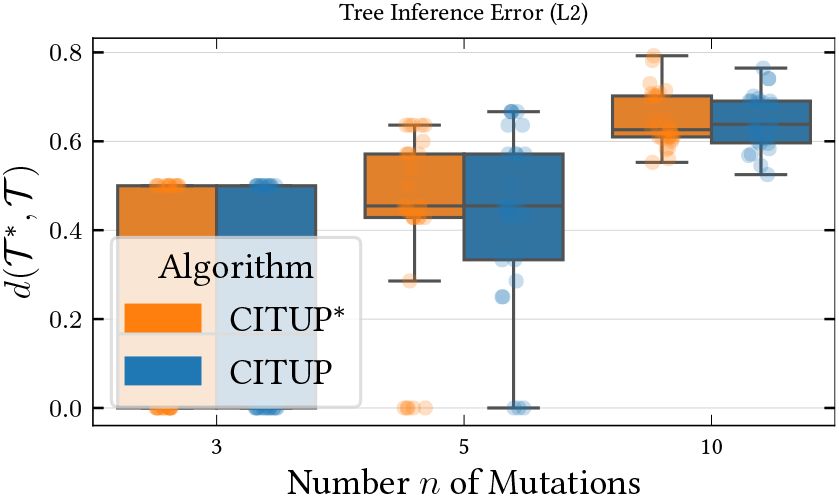
Ancestor-descendant distance between true and inferred phylogenies of CITUP [11] and CITUP^*^ across 180 simulated tumor phylogenies. CITUP^*^ is the result of replacing the CPLEX solver for the *ℓ*_2_ loss in CITUP with *fastppm*-L2.

**Fig. S6.**
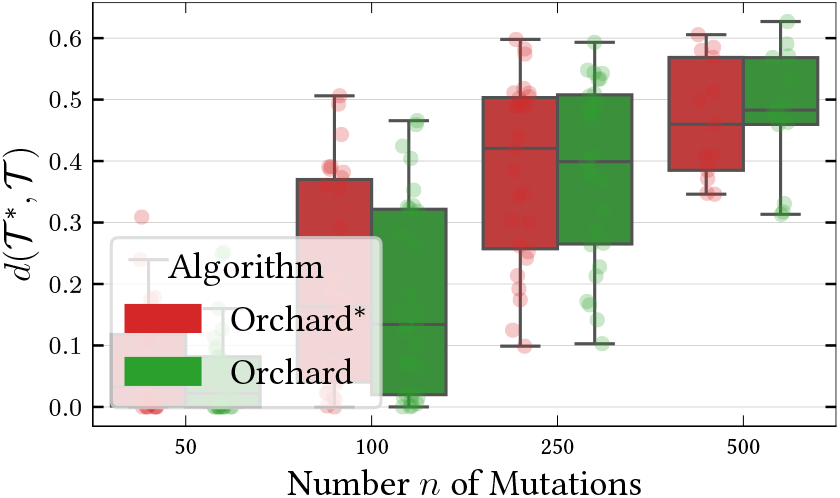
Ancestor-descendant distance between true and inferred phylogenies of Orchard [14] and Orchard^*^ across 120 simulated tumor phylogenies. Orchard^*^ is the result of replacing the projectppm solver for the *ℓ*_2_ loss in Orchard with *fastppm*-L2.

**Fig. S7.**
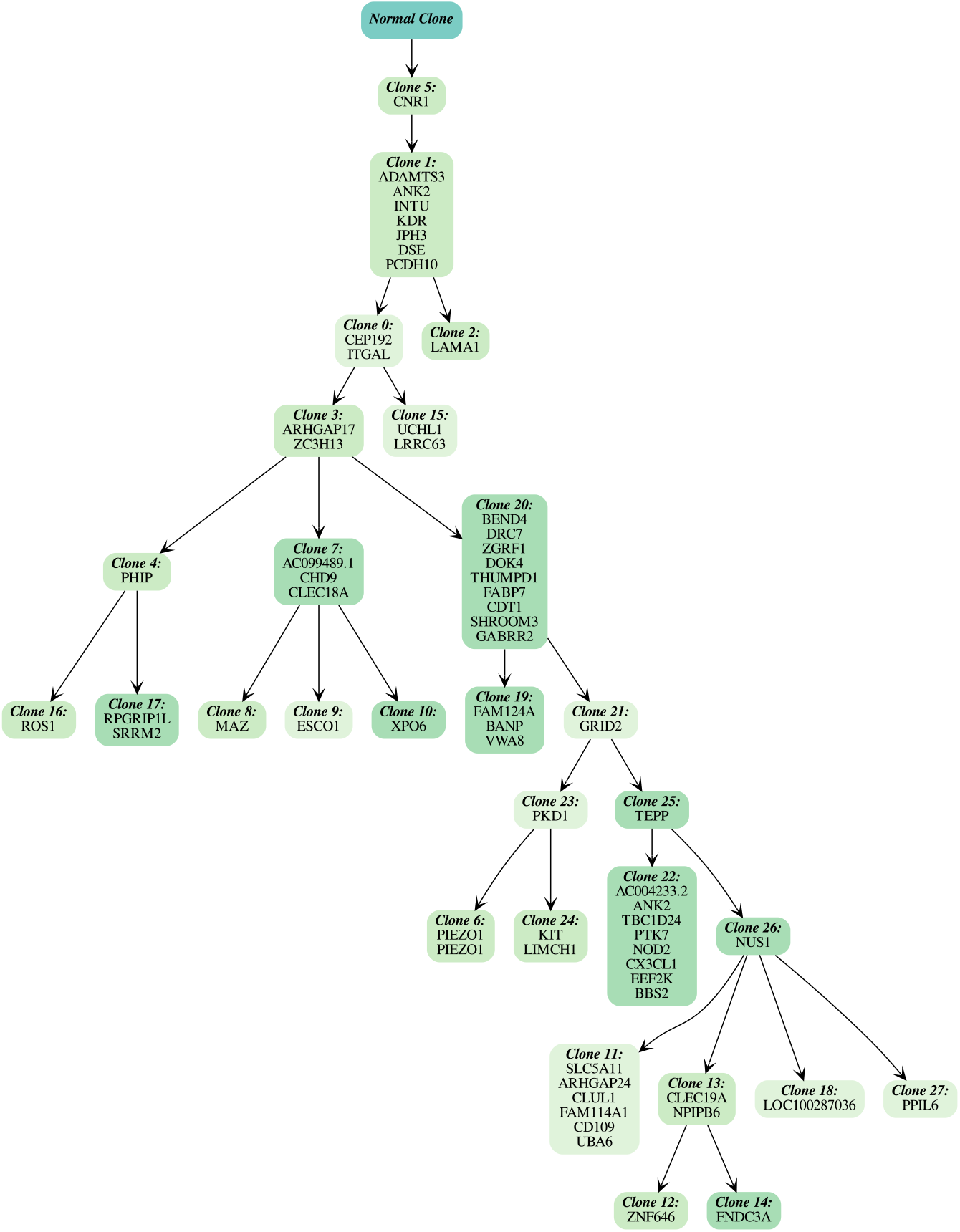
The phylogeny inferred by Orchard [14] on patient-derived xenograft POP66 [44]. For simplicity of visualization, degree-2 nodes are collapsed into clusters.

**Fig. S8.**
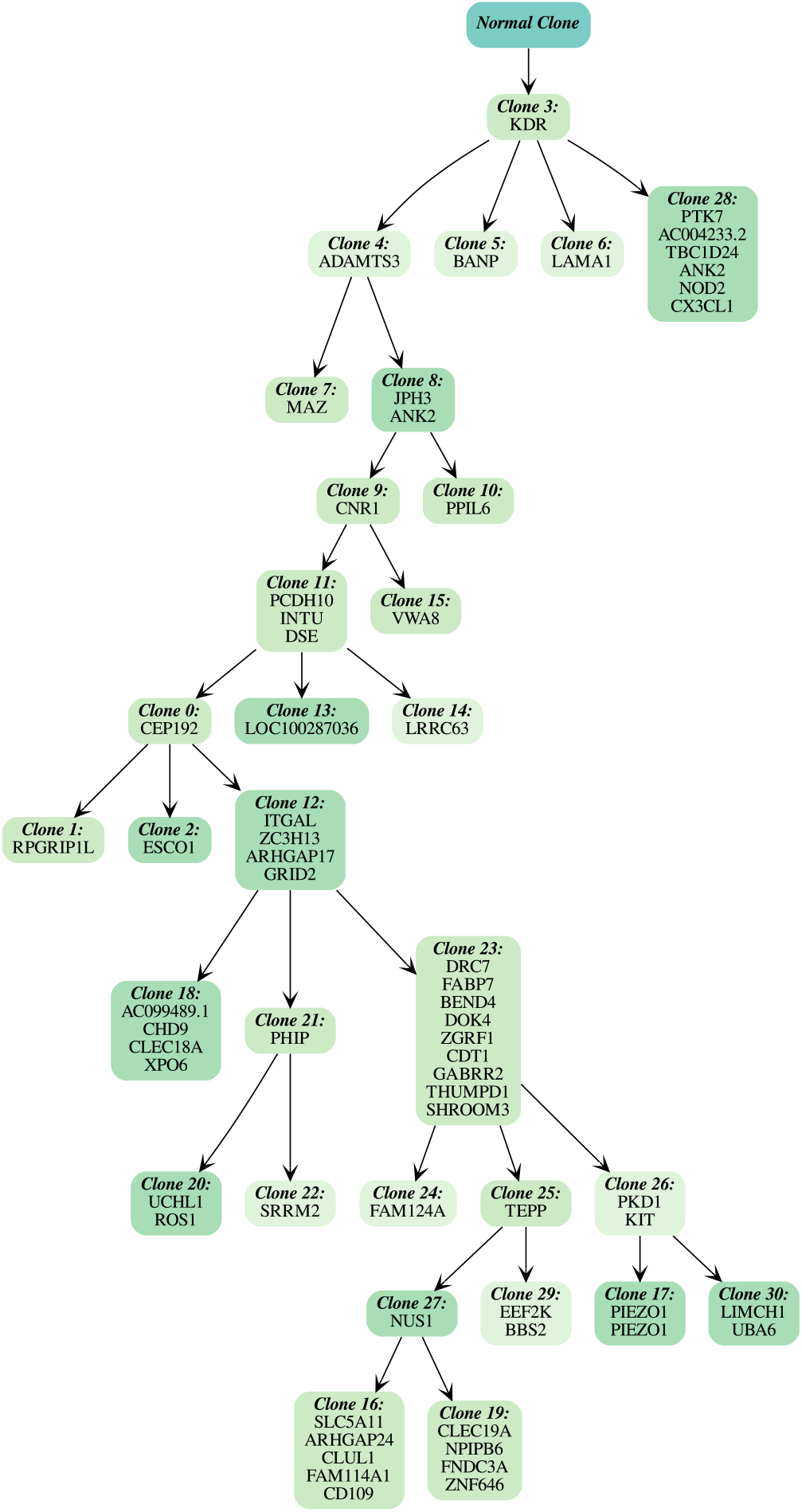
The phylogeny inferred by Sapling^*^ on patient-derived xenograft POP66 [44]. For simplicity of visualization, degree-2 nodes are collapsed into clusters.

## Footnotes

1 A mutation cluster *C* ⊆ [*n*] is summarized as an individual mutation *i* in one of two ways. First, one can pool reads, i.e. *vpi* ≜∑ *j∈ C vpj* and *dpi* ≜ ∑*j ∈C dpj*. Alternatively, to maintain a similar level of variance as the average mutation in the cluster, one can compute the average cluster 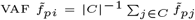 and set 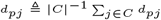 followed by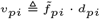, as done in [1].

## Notes

### Competing Interest Statement

The authors have declared no competing interest.

